# Comprehensive Molecular Characterization of Mitochondrial Genomes in Human Cancers

**DOI:** 10.1101/161356

**Authors:** Yuan Yuan, Young Seok Ju, Youngwook Kim, Jun Li, Yumeng Wang, Yang Yang, Inigo Martincorena, Chad J. Creighton, John N. Weinstein, Yanxun Xu, Leng Han, Hyung-Lae Kim, Hidewaki Nakagawa, Keunchil Park, Peter J. Campbell, Han Liang, PCAWG Network

**Affiliations:** Department of Bioinformatics and Computational Biology, The University of Texas MD Anderson Cancer Center, Houston, TX77030, USA; Cancer Genome Project, Wellcome Trust Sanger Institute, Hinxton,United Kingdom; Graduate School of Medical Science and Engineering, Korea Advanced Institute of Science and Technology, Daejeon 34141, Korea; Department of Health Science and Technology, Samsung Advanced Institute for Health Science and Technology, Sungkyunkwan University School of Medicine, Seoul, 135-710, Korea; Graduate Program in Structural and Computational Biology and Molecular Biophysics, Baylor College of Medicine, Houston,TX 77030, USA; Division of Biostatistics, The University of Texas Health Science Center at Houston, School of Public Health, Houston, Texas 77030, USA; Department of Medicine and Dan L. Duncan Cancer Center Division of Biostatistics, Baylor College of Medicine, Houston, TX 77030, USA; Department of Systems Biology, The University of Texas MD Anderson Cancer Center, Houston, TX 77030, USA; Department of Applied Mathematics and Statistics, Johns Hopkins University, Baltimore, MD21218, USA; Department of Biochemistry and Molecular Biology, The University of Texas Health Science Center at Houston McGovern Medical School, TX 77030, USA; Department of Biochemistry, Ewha Womans University School of Medicine, Seoul 03760, Korea; RIKEN Center for Integrative Medical Sciences, Tokyo 108-8639, Japan; Division of Hematology/Oncology, Innovative Cancer Medicine Institute, Samsung Medical Center, Seoul, 135-710, Korea; Cambridge University Hospitals NHS Foundation Trust, Cambridge, United Kingdom

## Abstract

Mitochondria are essential cellular organelles that play critical roles in cancer development. Through International Cancer Genome Consortium, we performed a multidimensional characterization of mitochondrial genomes using the whole-genome sequencing data of ~2,700 patients across 37 cancer types and related RNA-sequencing data. Our analysis presents the most definitive mutational landscape of mitochondrial genomes including a novel hypermutated case. We observe similar mutational signatures across cancer types, suggesting powerful endogenous mutational processes in mitochondria. Truncating mutations are remarkably enriched in kidney, colorectal and thyroid cancers and associated with the activation of critical signaling pathways. We find frequent somatic nuclear transfers of mitochondrial DNA (especially in skin and lung cancers), some of which disrupt therapeutic target genes (e.g., ERBB2). The mitochondrial DNA copy number shows great variations within and across cancers and correlates with clinical variables. Co-expression analysis highlights the function of mitochondrial genes in oxidative phosphorylation, DNA repair, and cell cycle; and reveals their connections with clinically actionable genes. Our study, including an open-access data portal, lays a foundation for understanding the interplays between the cancer mitochondrial and nuclear genomes and translating mitochondrial biology into clinical applications.

---

Mitochondria are crucial cellular organelles in eukaryotes, and there can be several hundred mitochondria in a single human cell^1^. Known as “the powerhouse of the cell”, mitochondria play essential roles in generating most of the cell’s energy through oxidative phosphorylation^2^. Despite its small size (16.5 kb), the circular mitochondrial genome includes 13 protein-coding genes that are equipped with all the elements necessary for their own protein synthesis^3^. The proteins encoded by mitochondrial DNA (mtDNA) genes work with other nuclear genes to form the respiratory chain complexes that are the main energy production system in cells. The involvement of mitochondria in carcinogenesis has long been suspected^4, 5^ because altered energy metabolism is a common feature of cancer^6^. Furthermore, mitochondria play important roles in other tasks, such as signaling, cellular differentiation, apoptosis, maintaining control of the cell cycle and cell growth^7^, all of which are intrinsically linked to tumorigenesis.

Several recent studies have performed the molecular characterization of mitochondria in cancer using next-generation sequencing data^8-12^, but these studies have usually described one specific dimension of the mitochondrial genome (e.g., somatic mutations) based on relatively small sample cohorts. Thus, a comprehensive, multidimensional molecular portrait of mitochondria across a broad range of cancer types has not been achieved. Furthermore, previous studies have focused on the patterns of mitochondrial alterations alone, without fully exploring the interplay between the mitochondrial genome and the nuclear genome, as well as the biomedical significance of mitochondrial alterations.

The International Cancer Genome Consortium (ICGC) project has deeply sequenced thousands of whole genomes from many cancer types, creating a tremendous resource for characterizing cancer mitochondrial genomes at an unprecedented level^13^. Meanwhile, The Cancer Genome Atlas (TCGA) project has generated RNA-seq data from a large number of patient samples, which allow assessing the transcriptional activities of mitochondrial genes^14^. Combining these two large-scale datasets, we first characterized the landscape of mitochondrial somatic mutations, nuclear transfers and copy numbers, and then examined the expression profiles of mitochondrial genes and their connections with clinically relevant nuclear genes. Finally, we developed an open-access data portal to facilitate the community-based investigation of these mitochondrial molecular data.

## Results

### The landscape of somatic mutations in cancer mitochondrial genomes

To characterize somatic mutations in mitochondrial genomes across cancer types, we extracted the mtDNA mapped reads of 2,658 cancer and matched normal sample pairs from the ICGC whole-genome sequencing (WGS) data. The samples we surveyed cover 21 cancer tissues and 37 specific cancer types (**Supplementary Table 1**). On average, the sequence depth for the mitochondrial genome was 9,959×, which was much higher than those obtained from the whole-exome sequencing data, allowing for confident detection of somatic mutations at a very low variant allele frequency (VAF; heteroplasmic) level (VAF>1% in this study; **Supplementary Fig. 1**). By applying a well-designed computational pipeline that carefully considered various potentially confounding factors (e.g., sample cross-contamination, mis-mapping of reads from nuclear mtDNA-like sequence^15^, and artifactual mutations caused by oxidative DNA damage during library preparation^16^), we identified a total of 7,611 substitutions and 930 small indels in 2,536 cancer samples (122 samples were excluded in the mutation analysis for the issues mentioned above; **Supplementary Fig. 2**, **Online Methods**). The somatic alterations called by our pipeline are highly reliable, as confirmed by a) comparison to a long-range PCR-based benchmark dataset (**Supplementary Table 2**) and b) the expected mitochondria-specific mutational signatures displayed by the low-allele-frequency variants (**Supplementary Fig. 3**). Across all the cancer samples, we observed several mutational hotspots in the regulatory D-loop region and the ND4 gene (**Fig. 1a**). As for the gene mutation frequency, *ND5* was the most frequently mutated gene in most cancer types, while *ND4* was the one in prostate and lung cancers, and *COX1* in breast, cervical and bladder cancers (**Fig. 1b**). We identified that cancer type and gene identity were the most important factors affecting the mutation status of the 13 coding genes (log-linear model, p [cancer type] < 2.2×10^-16^, p [gene] < 2.2×10^-16^) but the effect of their interaction was not significant (p [cancer type × gene] = 0.12). We also checked the expression of mtDNA somatic mutations using related RNA-seq data. Generally, VAFs of mtDNA mutations at the RNA level were consistent with those at the DNA level. However, a fraction of tRNA mutations showed significantly higher VAFs a^17^ (**Supplementary Fig. 4**).

**Figure 1.**
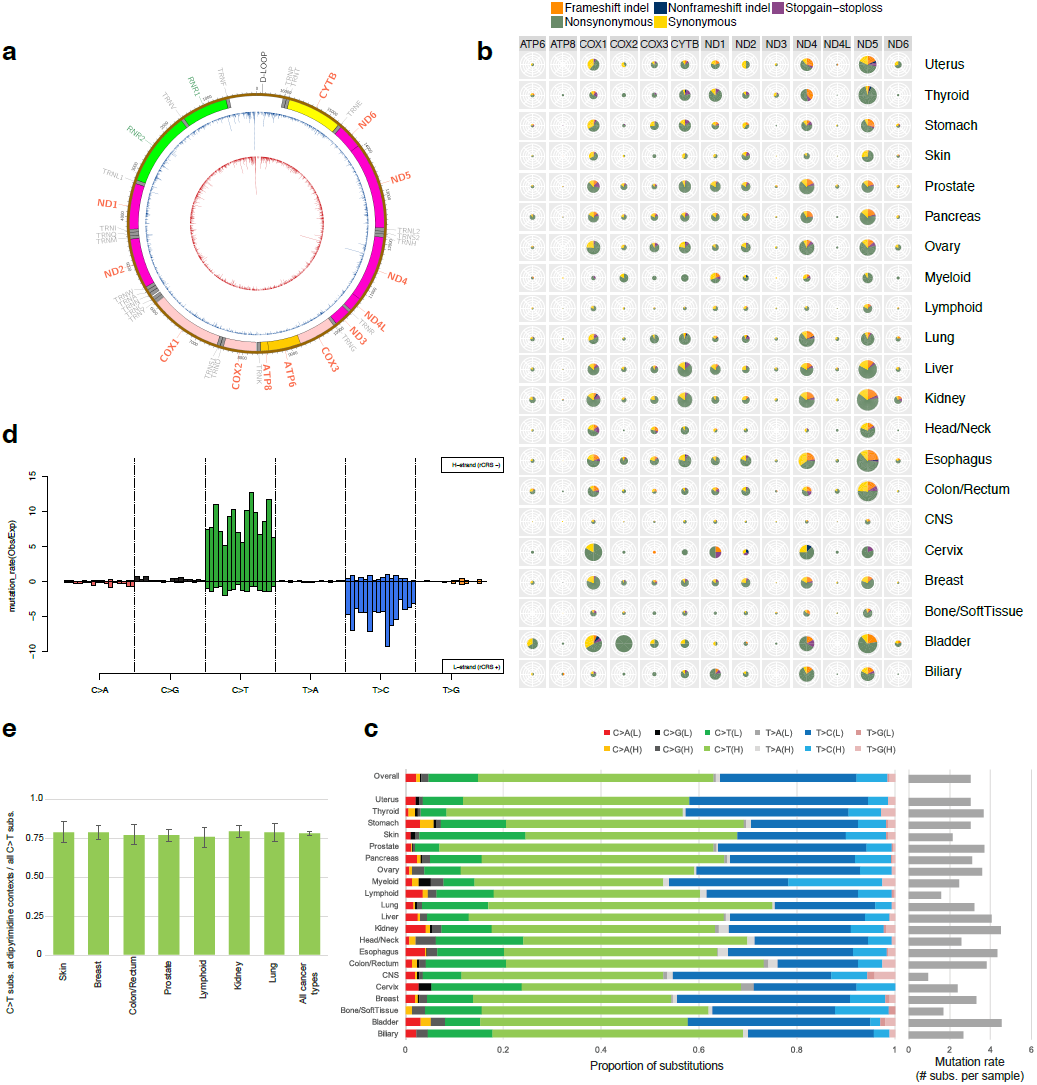
The mtDNA mutational landscape across different cancer tissues. (a) The landscape of mtDNA somatic substitutions identified, with the outer circle (blue) displaying all variants with variant allele frequency (VAF) > 1%, and the inner circle (red) displaying those with VAF >3%. (b) An overview of the mutation frequency of the 13 mtDNA coding genes across cancer tissue types. The size of each pie chart is proportional to the overall mutation frequency, with different color slices corresponding to different variant types. (c) Highly consistent mtDNA mutational signatures across 21 cancer tissue groups. The average numbers of somatic substitutions per sample are shown with bars on the right. (d) Replicative strand-specific mutational spectrum of mtDNA substitutions. The relative mutation frequencies (# of observed / # of expected) by 96 trinucleotide contexts are shown (top) for the H strand (bottom) for the L strand. (e) The proportion of C→T substitutions at dipyrimidine contexts (where UV light frequently generates mutations). No significant differences were observed between melanoma and other cancer types.

The mutational spectrum in the nuclear genomes shows highly distinct cancer-specific patterns^18^. For example, the mutational spectrum of lung cancers is dominated by C→A mutations associated with exposure to the polycyclic aromatic hydrocarbons in tobacco smoke^19^; while melanoma shows a distinct pattern of frequent C→T mutations at adjacent pyrimidines that is caused by the mis-repair of ultraviolet radiation-induced intrastrand crosslinks^20^. In contrast, although the mutation rate varied largely by cancer type, the mutation signatures in the mitochondrial genomes were very similar, with C→T (58.3%) and T→C (34.2%) substitutions being the most and second most frequent mutation types across all cancer types (**Fig. 1c, d**). Consistent with a previous report^10,^ we observed extreme replicational DNA strand bias, i.e., C→T substitutions occurring mostly on the mtDNA heavy (H) strand and T→C substitutions on the mtDNA light (L) strand (**Fig. 1d**) despite the relative depletion of cytosines and thymines on the mtDNA H and the L strand, respectively. Indeed, the fraction of C→A mutations (the dominant mutation type by tobacco smoking) in the mitochondrial genomes of lung cancer and the fraction of C→T mutations at dipyrimidine sites (the dominant mutation type by ultraviolet light) in melanoma were very similar to those in other cancers (**Fig. 1e**), suggesting that tobacco smoking and ultraviolet radiation are not the dominant causes of mtDNA mutations even in these two cancer types. The proportion of G→T mutations, which is the major mutation type induced by reactive oxygen species^21^, was very low (3.2%), suggesting that the impact of oxidative stress on mtDNA mutations is minimal contrary to the conventional wisdom. These observations indicate that the mutational processes in the mitochondrial genome are independent from those in the nuclear genome and that there is a strong unique endogenous mutational process operating in mitochondria.

On average, each sample had ~3 somatic substitution mutations in the mitochondrial genome (**Supplementary Fig. 5a**). Interestingly, some samples contained extremely large numbers of mtDNA mutations. The most striking case was a breast cancer sample harboring 33 mutations, 29 of which were localized in a 2kb region (**Fig. 2a**). The vast majority of these mutations were confirmed in the independent whole-exome sequencing data from the same patient (**Supplementary Fig. 5b, c**). These localized mutations showed highly similar VAFs (~7%), were mostly T→C substitutions (**Fig. 2b**) and were present on the same mitochondrial genome copies (co-clonal) when any pair of mutations were close enough to be phased by Illumina sequencing reads (i.e., < 500 bp). These suggest that all these mutations were acquired simultaneously by a single mutation crisis event (**Fig. 2c**), rather than by gradual accumulation of mutations over time. These features are quite similar to *Kataegis*, a pattern of localized hypermutation in nuclear genomes^22^. These mutations were enriched in the vicinity of mtDNA L-strand replication origin (**Fig. 2c**), suggesting that the event is coupled with the initiation of mtDNA L-strand replication.

**Figure 2.**
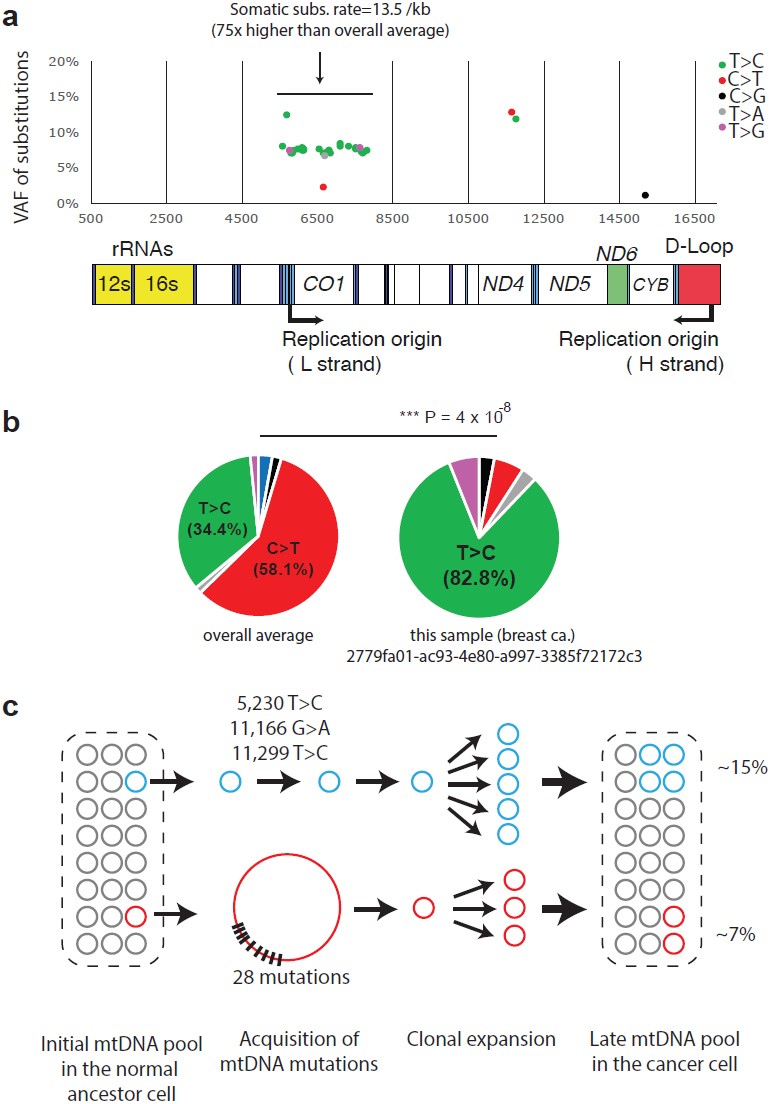
A hypermutated mitochondrial genome. (a) Somatic mutations are clustered in a 2kb region near the replication origin. (b) The T→C substitutions are dominant in this sample. (c) A proposed model of mutation acquisition.

### Excessive truncating mitochondrial mutations in kidney, colorectal and thyroid cancers

To investigate the mutational patterns of mtDNA genes, we examined the dN/dS ratio^10, 23^,a common measure of selective pressure. We found that dN/dS was close to 1 for missense mutations at different VAFs across cancer types, suggesting that overall selective pressure on mtDNA genes is nearly neutral (**Supplementary Fig. 6**). We next focused on truncating mutations and examined their heteroplasmic frequency distribution since they would lead to loss-of-function of individual mitochondrial genes. The frequency was remarkably low in most cancer types, suggesting the prevalence of negative selection; whereas kidney, colorectal and thyroid cancers showed substantially larger proportions of high-allele-frequency truncating mutations, suggesting the presence of positive selection in these cancer types (*F* test, *p* < 2.2×10^-16^, **Fig. 3a**). The enrichment of higher allele-frequency (i.e., >60% VAF) truncating mutations in these cancers, especially in kidney chromophobe and kidney papillary, was very striking, suggesting that loss of normal mitochondrial function is an important step in cellular transformation^24^ (**Fig. 3b**, **Supplementary Fig. 7)**. Interestingly, we did not observe such a high frequency of mtDNA truncating mutations in the samples of kidney clear cell, where inactivation of genes controlling cellular oxygen sensing (i.e. *VHL*) are observed in ~60% of cases^25^.

**Figure 3.**
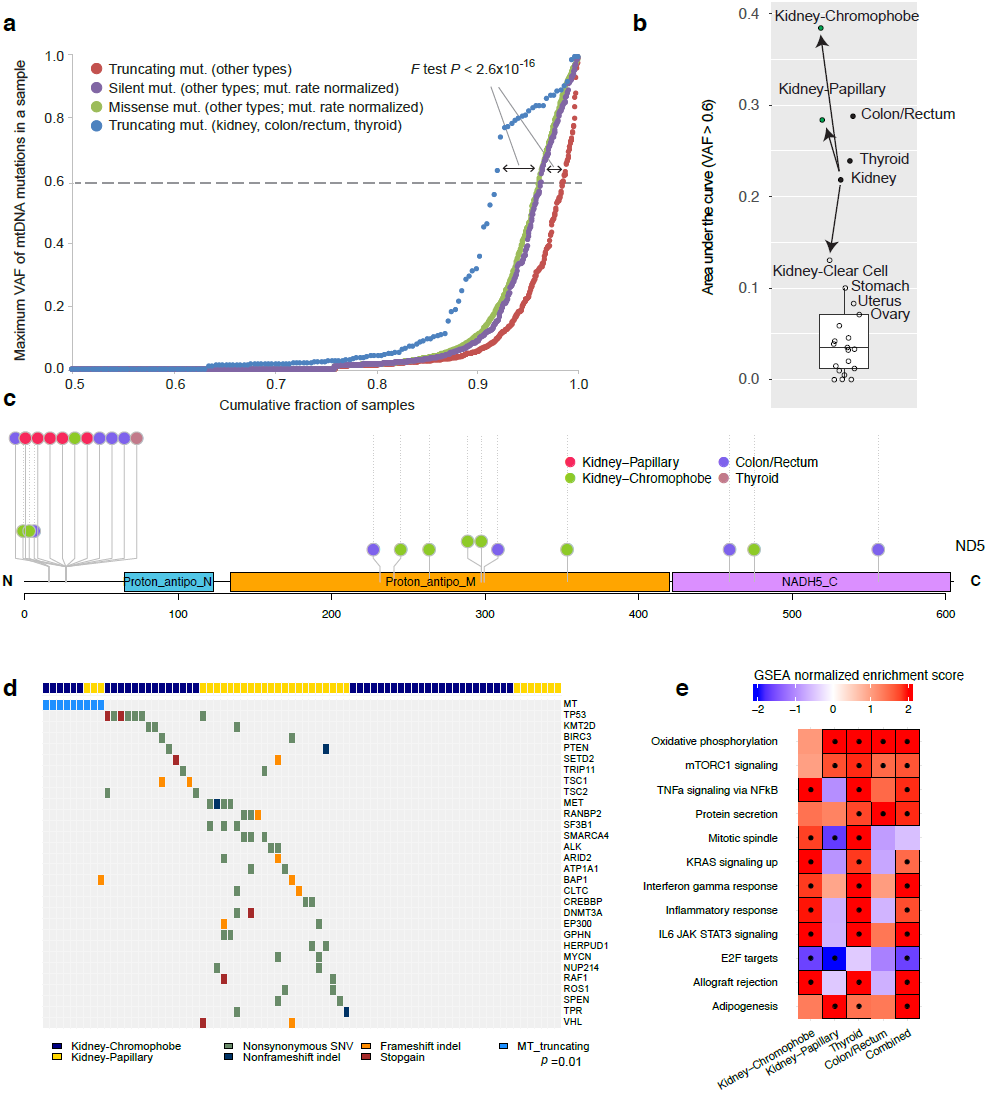
The mtDNA truncating mutation patterns. (a) The distinct VAF accumulation curves of truncating mutations between kidney/ colorectal/thyroid cancers and other cancer types. For comparison, similar curves were generated for silent and missense mutations in other cancer types (which are overall functionally neutral) after normalization of mutation numbers. Generally, fewer truncating mutations were observed in higher allele-frequency levels, presumably because those truncating mutations provide selective disadvantages to the mitochondria (or cells) that carry them. (b) Kidney chromophobe, kidney papillary, colorectal and thyroid cancers accumulated excessive high-allele-frequency (VAF >60%) truncating mutations. (c) Different distribution patterns of truncating mutations in ND5. (d) A heatmap representation of mtDNA truncating mutations with recurrent somatic mutations in cancegene in kidney chromophobe and kidney papillary. (e) A heatmap representation of signaling pathways enriched by nuclear genes upregulated in in cancer samples with truncating mutations. A dot indicates FDR < 0.05

We then examined the truncating mutations in *ND5*. This mtDNA gene showed the highest mutation rate across cancer types, and in particular, it significantly enriched truncating mutations in kidney chromophobe (chi-squared test, *p* < 6×10^-3^). Interestingly, we observed distinct distribution patterns of truncating mutations in different tumor contexts: more enriched at N-terminal for kidney papillary but more dispersed for kidney chromophobe and colorectal cancer (Fisher exact test, *p* = 0.05, **Fig. 3c**). Integrating with the mutation data of nuclear genes, we found that the high-allele-frequency truncating mutations in the two kidney cancers were mutually exclusive to the mutations of known cancer genes (Fisher exact test, *p* = 0.01, **Fig. 3d**). Moreover, gene set enrichment analysis (GSEA)^26^ showed that up-regulated genes in cancer samples with truncating mutations were significantly enriched in critical pathways such as oxidative phosphorylation, mTOR signaling, TNFα signaling, protein secretion, suggesting activation of these pathways (false discovery rate [FDR] < 0.05, **Online Methods, Fig. 3e**). These results suggest functional roles of mitochondrial truncating mutations in the initiation and/or development of kidney, colorectal and thyroid cancers.

### Somatic transfer of mitochondrial DNA into the nuclear genome

Recently, somatic transfers of mtDNA into the nuclear genome have been reported^11^, mostly in breast cancer. In order to understand the general pattern of somatic mtDNA nuclear transfers (SMNTs) in human cancer, we applied the same bioinformatic pipeline to the 2,658 cancer genomes. In total, we found 55 positive cases (2.1% overall positive rate) (**Online Methods**). However, the frequency of SMNT varied according to the cancer tissue type (Fisher’s exact test, *p* < 1×10^-5^, **Fig. 4a**). In particular, samples from lung, skin, breast, and uterine cancers showed frequencies higher than 5%. Among lung cancers, lung squamous showed 14.6% positive rate (7/48), a significantly higher rate than the average (Fisher’s exact test, *p* < 0.001). Despite the overall prevalence of 2%, we did not find any positive cases from >600 samples of blood, kidney, esophagogastric, liver, prostate and colorectal cancers. This tissue-specific pattern suggests that the processing of mtDNA in cells, from mitochondria to integration into nuclear genomes, depends on specific cancer contexts. We further examined the correlations of SMNT with structural variations in the nuclear genome. Intriguingly, across cancer types, the samples with SMNT showed a much higher number of global and local structural variations in the nuclear genome than those without (*p* = 1×10^-4^; **Fig. 4b**), and this pattern was particularly obvious in breast cancer (*p*= 5.4×10^-4^). Further, this pattern was also consistent across different types of structural variations (**Supplementary Fig. 8**) Indeed, the distance from SMNT breakpoints to the nearest structural variation breakpoints was significantly shorter than random expectation, especially for inversions and translocations, but not for deletions and tandem duplications (**Fig. 4c**). These results suggest that the integration of mtDNA segment into nuclear DNA is often mechanistically combined with some specific mutational processes of structural variations.

**Figure 4.**
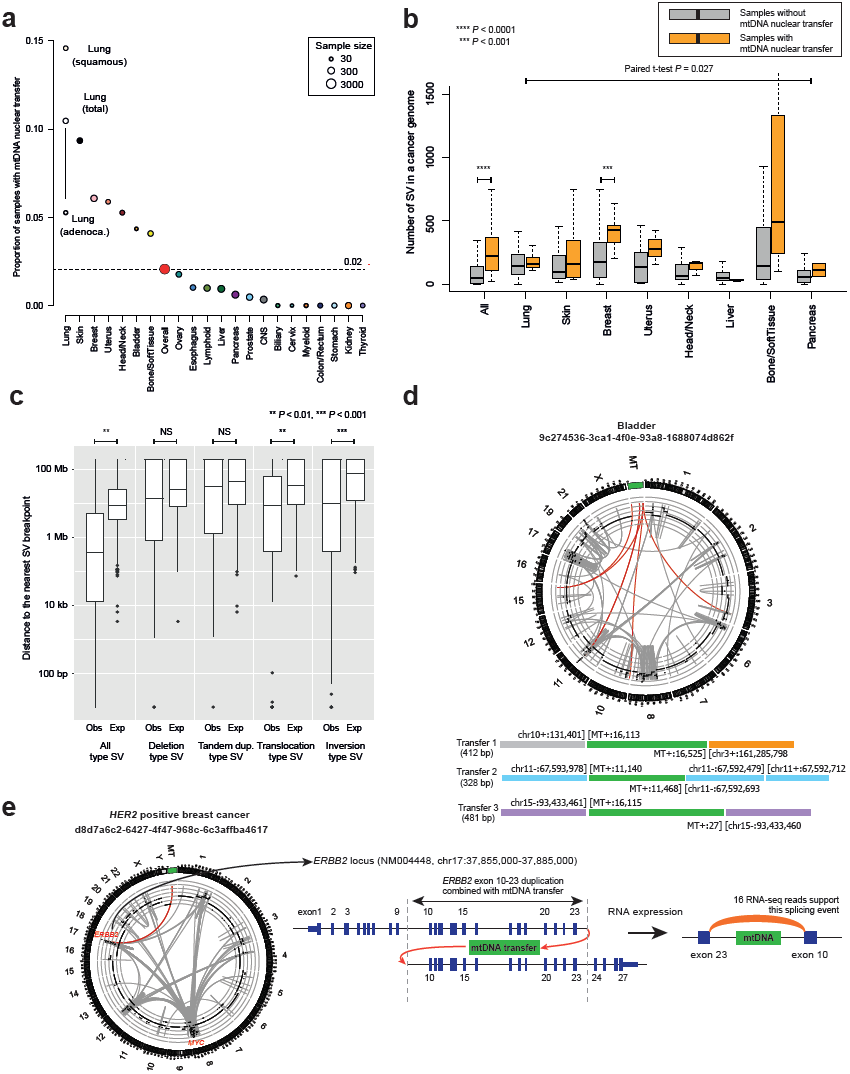
Somatic transfer of mitochondrial DNA into the cancer nuclear genome. (a) The frequency of SMNT in different cancer tissues. (b) The numbers of SV breakpoints in samples with and without SMNT. (c) Distance from SMNT breakpoints to the nearest SV breakpoints are shorter than random expectation for all and each type of SVs. (d) A Circos plot representing three independent somatic mtDNA nuclear transfer events in a bladder cancer genome. Twenty-three human chromosomes are shown in the outer layer. Copy numbers of nuclear cancer genomes are represented by black dots in the inner layer. Chromosomal rearrangements are shown with gray curves and mtDNA nuclear transfers are represented by red curves. The summary of three nuclear transfers is depicted below the Circos plot with breakpoints. (e) A mtDNA nuclear transfer found in a HER2-positive breast cancer genome, leading to a tandem duplication process of ERBB2 exons 10–23. The novel exon junction is supported by the RNA reads from the corresponding RNA-seq data of the patient sample.

Despite the overall low SMNT frequency (~2%), some cancer samples showed multiple independent SMNT events (**Fig. 4d, Supplementary Fig. 9**). For example, a bladder cancer genome included three small mtDNA segments (all < 500 bp, **Fig. 4d**); and a breast cancer genome contained two independently transferred mtDNAs, with one transferred mtDNA being almost the entire mtDNA genome **(Supplementary Fig. 9a)**. We observed complex rearrangements even in the mtDNA segments somatically transferred to the nucleus (**Supplementary Fig. 9b**), implying extreme genome instability during the SMNT process. Furthermore, SMNTs in 35 cases showed mtDNA insertion in the middle of genes (n = 42), mostly in introns (n = 37) but also in protein-coding regions (n = 3) and in untranslated-regions (n = 2) (**Supplementary Table 3**). Among these, open reading frames of at least 23 genes (23/42 = 55%), including cancer genes such as *ERBB2, FOLH1* and *ULK2*, were predicted to be altered by these SMNTs and their combined SV events in the vicinity (**Supplementary Fig. 10**). Of particular interest, one SMNT was integrated into the *ERBB2* gene in a HER2-positive breast cancer genome (**Fig. 4e**), combined with frame-shifted duplication events of exons 10–23. This example suggests that some SMNTs influence functionally and clinically important chromosomal rearrangements cancer cells.

### Mitochondrial copy number alterations across cancer types

Although previous studies have examined mtDNA copy numbers in individual cancer types^27-29^ or from a collection of whole-exome sequencing data^12^, we performed a systematic analysis of mtDNA copy numbers over the largest sample cohort with WGS data so far. The analyses of mtDNA copy numbers are usually based on the ratio of the coverage depth of mtDNA over that of nuclear DNA. However, since (a) cancer samples consist of both normal and cancer cells (i.e., tumor purity < 1) and (b) cancer nuclear genomes have frequently gone through somatic copy number alterations30 (i.e., ploidy ≠ 2), the simple estimation of mtDNA copy number without accounting for these factors could be biased. Therefore, we incorporated tumor purity and ploidy data in our estimation of mtDNA copy numbers (**Online Methods**). Compared with the simple estimation, our method can effectively correct for the bias introduced by the low purity and high ploidy of cancer samples^12^ (**Supplementary Fig. 11**).

Based on the 2,157 cancer samples that passed the purity filter (**Online Methods**), we observed a great variation in mtDNA copy numbers among and within cancer tissues: mtDNAs were most abundant in samples of ovarian cancer (median, 644 copies per cell) and least abundant in myeloid cancer (median, 90 copies per cell) (**Fig. 5a**). Furthermore, the cancer types that were derived from the same tissue frequently showed distinct mtDNA copy number distributions (**Fig. 5b**, **Supplementary Fig. 12**). For example, the mtDNA copy numbers of kidney chromophobe were significantly higher than those of kidney clear cell and kidney papillary (ANOVA, *p* < 7.8×10^-6^). This may be interlinked with the general inadequacy of mitochondrial quality control and resultant increase of steady-state mtDNA copy number, as seen in renal oncocytoma^42^. Indeed, we found that the mtDNA copy number was significantly higher in the samples with high-allele-frequency truncating mutations (ANOVA, *p* < 1.7×10^-4^, **Fig. 5c**), suggesting that the dosage effect of mtDNAs was selected to compensate for the deleterious effect of truncating mutations. For the cancer samples with the WGS data from matched normal tissues, we observed increased mtDNA copy numbers in cancer samples in patients with chronic lymphocytic leukemia, lung squamous and pancreatic adenocarcinoma, but decreased copy numbers in cancer samples in patients with kidney clear cell, liver hepatocellular carcinoma and myeloproliferative neoplasm (**Fig. 5d**). The distinct pattern in different cancer types may be due to cancer-specific oncogenic stimulation, metabolic activity and mitochondrial malfunctions. For example, a recent study^12^ suggests that significantly decreased mtDNA copy number in kidney clear cell may be due to down-regulation of PGC-1α, a central regulator of mitochondrial biogenesis by hyperactivated HIF-1α, which is most frequently mutated and activated in this cancer type^31^.

**Figure 5.**
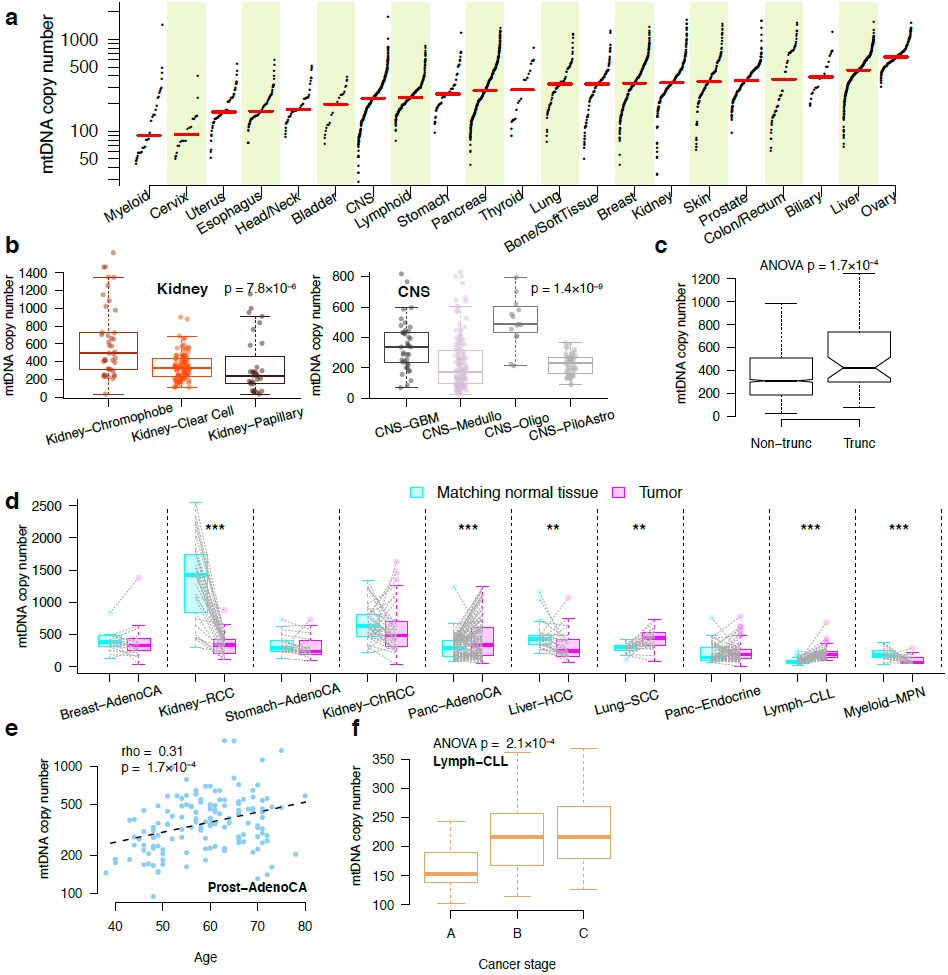
A pan-cancer view of mtDNA copy number. (a) The distributions of mtDNA copy number by cancer tissue type. (b) Distinct mtDNA copy number distributions for cancer types derived from kidney and brain. (c) mtDNA copy numbers with and without truncating mutations in mtDNA genes. (d) Paired copy number comparison in tumor samples and their matching normal tissue samples. (e) Correlation of mtDNA copy numbers with the patient age in prostate cancer. (f) Correlation of mtDNA copy number with patient stage in chronic lymphocytic leukemia.

To assess the biomedical significance of mtDNA copy numbers, we examined their correlations with key clinical variables. We found significant positive correlations between the mtDNA copy number and the patient’s age in prostate (Spearman rank rho = 0.31, *p* < 1.7×10^-4^, **Fig. 5e**), colorectal and skin cancers (**Supplementary Fig. 13**). In contrast, we observed negative correlations of normal blood mtDNA copy number with patient age in most cases (**Supplementary Fig. 14**). Moreover, we observed the correlations between mtDNA copy number and tumor stage in multiple cancer types (**Fig. 5f**, **Supplementary Fig. 15**).

### The co-expression network analysis of mitochondrial genes

We quantified the expression levels of the 13 mtDNA genes using RNA-seq data profiled from 4,689 TCGA tumor samples of 13 cancer types (**Supplementary Table 4**). The correlation between the expression levels of the mtRNA genes and mtDNA copy number varied by cancer types (**Supplementary Fig. 16**). Among cancer types, the mtDNA genes were highly expressed in the three types of kidney cancer (chromophobe, papillary and clear cell) but lowly expressed in the three types of squamous cell carcinoma (cervical, lung, and head&neck) (**Fig. 6a**). This observation was partially due to the relative abundance of mtDNA copy number across cancer types; and it is also consistent with a study of normal tissues^32^.

**Figure 6.**
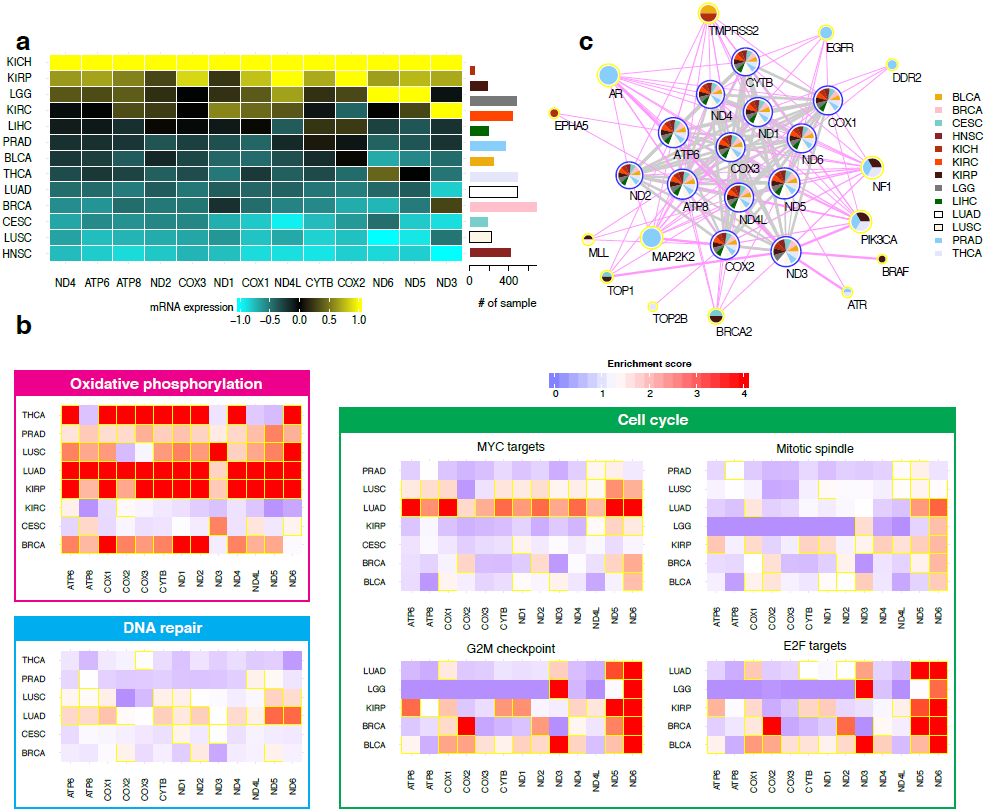
Co-expression patterns of mtDNA gene across different cancer types. (a) An overview of the expression levels of the 13 mtDNA genes of 13 cancer types in a heatmap (left) and the sample sizes in a barplot (right). (b) The commonly enriched pathways identified by the co-expression with mtDNA genes in different cancer types. The borders of the cells with FDR < 0.05 are highlighted in yellow. (c) An mtDNA gene-centric pan-cancer co-expression network. The colors of the pie chart at each node indicate the occurrences of the node in corresponding cancer types. The size of the node is proportional to the number of the direct neighbors (connectivity) of the node. The thickness of the edge is proportional to the frequency of this edge being observed across all cancer types. Further, the edges are colored according to the connection type: mtDNA gene–mtDNA gene connection in gray, mtDNA gene–nuclear gene connection in magenta

To gain more insight into the functions of mtDNA genes and their related nuclear genes and pathways, for each cancer type, we used the WGCNA package^33^ to build a weighted gene co-expression network that consisted of both nuclear genes and mitochondrial genes (**Online Methods**). We then performed GSEA^26^ on the basis of the rank of all nuclear genes by measuring their edge strength to a mitochondrial gene in the co-expression network. We found oxidative phosphorylation to be the top-ranked enriched pathway, and to be enriched in 8 out of the 13 cancer types examined (FDR < 0.05), highlighting the essential role of mitochondria in energy generation (**Fig. 6b**). Pathways related to the cell cycle (MYC targets, mitotic spindle, G2M checkpoint and E2F targets) and DNA repair were also enriched in multiple cancer types (**Fig. 6b**), consistent with the established notion that mtDNA plays an important role in these pathways^34, 35^

We also examined the mtDNA-centric co-expression networks (**Figure 6c**, **Online Methods**). Across cancer types, the mtDNA genes were almost always strongly interconnected, which is expected since they are transcribed as long polycistronic precursor transcripts^36^. Interestingly, several clinically actionable genes were among the neighboring genes that showed strong co-expression patterns with mtDNA genes (**Fig. 6c**, **Supplementary Figure 17**). For example, *AR*, *EGFR*, *DDR2* and *MAP2K*2 were connected with mtDNA genes in prostate cancer; and *TMPRSS2*, *NF1*, *PIK3CA*, *BRCA1* and *TOP1* were the top neighbors of mtDNA genes in multiple cancer types. These interactions highlight the clinical relevance of mtDNA genes, and elucidating the underlying mechanisms will lay a foundation for developing mtDNA-related cancer therapy.

### The Cancer Mitochondrial Atlas data portal

To facilitate mitochondria-related biological discoveries and clinical applications, we developed an open-access, user-friendly data portal “The Cancer Mitochondrial Atlas” (TCMA) for fluent exploration of the various types of molecular data characterized in this study (**Supplementary Fig. 18**). The data portal can be accessed at http://bioinformatics.mdanderson.org/main/TCMA:Overview. There are four modules in TCMA: somatic mutations, nuclear transfer, copy number, and gene expression. The first three modules are based on the ICGC WGS data and provide detailed annotations for the corresponding features of each cancer sample. The last module is based on the TCGA RNA-seq data and provides an interactive interface through which users can visualize the co-expression network with convenient navigation and zoom features. Users can not only browse and query the molecular data by cancer type, but also download all the data for their own analysis.

## Discussion

This work represents the first study to characterize the cancer mitochondrial genome in a comprehensive manner, including somatic mutations, nuclear transfer, copy number and mtDNA gene expression. Because of the mtDNA ultra-high coverage from the WGS data and the large number of patient samples surveyed, our study provides the most definitive landscape of mtDNA somatic mutations and reveals several unique features. First, in contrast to the diversified mutational signatures observed in the nuclear genomes of different cancers^18^, mtDNAs show very similar mutational signatures regardless of cancer tissue origins: predominantly C→T substitutions in the H strand and T→C substitutions in the L strand. This monotonous pattern may partially stem from different mutational generators implemented in mitochondrial DNA polymerase and/or differential DNA repair processes. operating between nucleus and mitochondria^10,37,38^. Due to their large numbers of copies pe cell, mitochondria may simply remove defective DNA through autophagy and other mitochondrial dynamic mechanisms^39^, rather than employing a complex array of repair proteins. Second, although hypermutated nuclear genomes have been reported in several cancer types (e.g., endometrial and colorectal cancers^40, 41^), our study here reports the first hypermuated mitochondrial case, highlighting the mutational complexity even in this tiny genome. Third, our systemic analysis of mitochondrial genomes have firmly demonstrated that several cancer types are enriched for high-allele-frequency truncating mutations, including previously reported kidney chromophobe^24, 42^ as well as newly identified kidney papillary, thyroid, and colorectal cancers. Interestingly, thyroid and kidney are the most frequent sites of oncocytoma, which are rare, benign tumors characterized by the vast accumulation of defective mitochondria due to pathogenic mitochondrial mutations^43^. Thus, some cancers with defective mitochondria may have grown from early benign oncocytic cells^42^.

One novel aspect of our study is the integrative analysis of mitochondrial molecular alterations with those in the nuclear genome that are characterized by ICGC. We found that high-allele-frequency truncating mtDNA mutations are mutually exclusive to mutated cancer genes in kidney cancer; mtDNA nuclear transfers are associated with increased numbers of structural variations in the nuclear genome; and the mtDNA co-expressed nuclear genes are enriched in several processes critical for tumor developments. These results indicate that the mitochondrial genome is an essential component in understanding the complex molecular patterns observed in cancer genomes and helping pinpoint cancer driver events. Importantly, our results, such as the nuclear transfer of mtDNA into a therapeutic target gene, correlations of mtDNA copy numbers with clinical variables, and the co-expression of mtDNA and clinically actionable genes, underscores the clinical importance of mitochondria.

Taken together, this study has untangled and characterized the full spectrum of molecular alterations of mitochondria in human cancers. Our analyses have provided essentially complete catalogs of somatic mtDNA alterations in cancers, including substitutions, indels, copy number and structural variations. Furthermore, we have developed a user-friendly web resource to enable the broader biomedical community to capitalize on our results. These efforts would lay a foundation for translating mitochondrial biology into clinical investigations.

## Online Methods

### Data generation and collection

We extracted BAM files of mitochondrial DNA sequencing reads from the whole-genome alignment files of 2,658 cancer samples and their matched normal tissue samples generated by the ICGC PanCancer Analysis of Whole Genomes group (PCAWG). BWA was used to align the reads to the human reference genome (GRCh37). From CGHub, we obtained the TCGA RNA-Seq BAM files of 13 cancer types, all of which employed paired-end sequencing strategies. We used Cufflinks to quantify the mRNA expression levels (in fragments per kilobase per million mapped fragments [FPKM]) of the 13 mitochondrial protein-coding genes.

### Somatic mutation calling

The nuclear genome mutations were called by using the Sanger pipeline, provided by the PCAWG. The mitochondrial varwere initially called using VarScan2^44^ aniants d the same parameter setting as previously reported^10^: --strand-filter 1 (mismatches should be reported by both forward and reverse reads), --min-var-freq 0.01 (minimum VAF 1%), --min-avgqual 20 (minimum base quality 20), --min-coverage × and --min-reads2 ×). We applied a series of downstream bioinformatic filters to further remove false positives as follows (**Supplementary Fig. 2a**).

First, we filtered germline polymorphisms and false positive calls. For example, frequent mapping errors due to known mtDNA homopolymers, candidates with substantial mapping strand bias, and candidates with substantial mutant alleles in the matched normal sample. For simplicity in the analyses, we removed multi-allelic mtDNA mutations and back mutations from non-reference to reference allele. After this filtration step, we obtained 10,083 somatic substitution candidates.

Second, we examined DNA cross-contamination, because even minor DNA cross-contamination (i.e., contamination level < 3%) would generate many low VAF false positive calls that are in fact germline polymorphisms from the contaminating sample. We tested whether mtDNA somatic mutations detected from a cancer sample show greater overlap with known mtDNA polymorphisms than expected from the overall average rate (73.5%; 3,922/5,337 substitutions) using the binomial test with cutoff *p* < 0.01. From this step, we removed 96 samples with evidence of DNA cross-contamination (harboured 935 known mutations out of 1,131 known mutation candidates).

Third, we examined the overall mtDNA substitution signatures in the 96 possible mutation classes. We removed four samples with extremely high proportions of C→G substitutions with strong sequence context bias (at CpCpN→CpApN; most frequently at CpCpG→CpApG; **Supplementary Fig. 2b**), which is known to be a spectrum from artificial guanine oxidation during sequencing library preparation steps^16^ with low VAF (1%–2%). We explicitly removed these samples from further analyses.

Then, we examined the possibility of false positive calls due to mis-mapping of reads from inherited nuclear mtDNA-like sequences (known as numts) not represented in the human reference genome^15,^ especially when the specific numts regions were amplified in the cancer nuclear genome. These mutation candidates showed some specific features: (1) appeared as highly recurrent mtDNA somatic mutations among multiple samples; (2) VAFs in mitochondria were only slightly higher than our 1% cutoff criteria; and (3) the matched normal samples also had small, but substantial numbers of mutation allele counts. To remove these false positive calls, we applied two statistical tests: (1) whether the VAF of a mutation candidate in the matched normal sequences was within the normal range (< 0.0024, the cutoff is determined by the median VAF of all mutation candidates + 2×IQR), and (2) whether

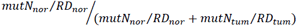

was within the normal range (<0.0357, the cutoff is determined by the median VAF of all mutation candidates + 2×IQR), where mutN is the mutation allele count, RD is the average read depth for the nuclear genome, and *nor* and *tum* are normal and matched tumor tissues, respectively. When a mutation appeared to be an outlier according to both criteria, we removed the candidate from our downstream analyses.

In our previous study^10^, we could not detect mutations under a 3% VAF cutoff because mtDNA were sequenced with a read depth of ~100× from the majority of the samples surveyed. Taking advantage of the ultra-high depth (>8,000×) in this study, we used a 1% VAF cutoff to obtain better sensitivity. We found 2,133 more substitutions when the VAF was between 1% and 3%. Because of the ultra-high depth, even 1% VAF mutations were considered to be specific, and were supported by a high number (n = ~80) of mutation alleles. We confirmed the high specificity of these mutations using the unique mtDNA mutational signatures robustly observed even from these low VAF mutations: (1) the mutational spectrum is generally consistent with those from higher heteroplasmic levels of mutations (i.e., VAFs from 3%-10%, 10%-100%); (2) we observed the absolute dominance of C→T ant T→C substitutions in the expected trinucleotide contexts (NpCpG for C→T and NpTpC for T→C substitutions); and (3) we also observed extreme replication strand bias (**Supplementary Fig. 3**). These features would not be observed if contaminations resulted in many false positive calls. To assess the factors affecting the mutation frequency of the 13 coding genes, we performed the sample-level analysis using log-linear modeling: we assigned the binary mutation indicator (1: with mutation; 0: without mutation) to each sample for each gene and then fit this binary response variable to a logistic regression model including cancer type, gene identity, and their interaction as explanatory variables, which were later summarized using analysis of variance (ANOVA).

### Truncating mutation analysis

Taking into account the mtDNA specific mutational signature, we examined the dN/dS ratio for mtDNA missense substitutions as reported previously^10^. We defined truncating mutation as those that lead to truncated protein products (i.e., nonsense mutations, frame-shift indels); and accordingly categorized the samples into the truncating group (bearing at least one truncating mutation with VAF ≥ 60%). The ND5 protein domain information was obtained from Pfam (http://pfam.xfam.org/protein/P03915). The list of cancer gene census was obtained from http://cancer.sanger.ac.uk/cosmic/download. Cancer census genes with recurrent somatic mutations in kidney chromophobe and kidney papillary were selected for the analysis of mutual exclusivity and heatmap representation. One sample with nuclear DNA hypermutator phenotype was excluded from this analysis. To examine the functional consequences of mtDNA truncating mutations, we performed GSEA based on the ranks of differentially expressed genes between samples with mtDNA truncating mutations and without for kidney chromophobe, kidney papillary, colorectal, and thyroid cancers and their combination, and identified significantly enriched pathways at FDR = 0.05.

### Somatic mtDNA nuclear transfer

We examined the whole-genome sequencing data from the cancer and matched normal tissue samples using a pipeline for the identification of mtDNA translocation to the nuclear genome as reported previously^11^The specificity was shown to be 100% in the previous study^11^Briefly, we extracted and clustered discordant reads from cancer genomes, where one end aligned to nuclear DNA and the other to mtDNA. Then, in order to determine the nucleotide resolution breakpoints, we searched for split reads near putative breakpoint junctions (1,000 bp upstream and downstream), where a fraction of a single read aligned to genomic DNA near the junctions and the rest aligned to mtDNA. All filtering criteria were the same as previously reported, except that we did not use BLAT^45^ for split-read detection because the BWA-MEM alignment tool used to map all pan-cancer samples fundamentally enables split-read mapping. We removed candidate mitochondria–nuclear DNA junctions that overlapped with clusters from matched and unmatched normal samples and/or known human SMNTs, a combined set from the human reference genome (hg19; n= 123) and from a published study^46^ (n = 766) because the source of the mtDNA sequence fused to the nuclear genome might be SMNTs rather than real mitochondria in the cytoplasm of cells. We obtained the structural variations from ICGC PCAWG and compared the samples with and without SMNT using t-test. To study the relationship of SMNT and SV breakpoints, we randomly chose the same number of SV breakpoints from each sample for 100 times to estimate the random expectation.

### MtDNA copy number analysis

To better estimate the mtDNA copy number (CN) for cancer samples, we employed the following formula that incorporates both tumor purity and ploidy information:

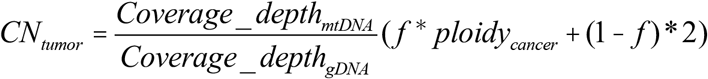, where *f* is the tumor tumor purity (ranging from 0 to 1, where 1 stands for pure cancer cells and 0 stands for pure normal cells); *Coverage* _*depth*_*mtDNA*_ and *Coverage* _ *depth*_*gDNA*_ are the mean coverage depths for mtDNA and the nuclear genome in individual WGS BAM files, respectively;*Ploidy*_*cancer*_ is the number of sets of chromosomes in tumor cells, while the ploidy in the normal cellsis 2. Both*f* and*Ploidy*_*cancer*_ were obtained by the allele-specific copy number analysis of tumors (ASCAT) estimation^47^, provided by PCAWG. Donors with multiple samples were pre-selected so that each donor comes with one representative primary cancer sample. We excluded cancer samples with low purity (<0.4, estimated by ASCAT) for further downstream analyses. Analysis of variance (ANOVA, if more than two cancer types) or t-test was used to compare the mtDNA copy number of cancer types derived from the same tissue. Since many of the normal samples are from blood, we focused on the cancer types with at least 10 samples from the normal tissue adjacent to the tumor in order to compare the mtDNA copy number of the paired cancer and normal samples. We used the Wilcoxon signed rank test to compare the mtDNA copy number for each selected cancer type and further adjusted the raw *p*-values by the FDR. To assess the correlation of mtDNA copy number with truncating mutations, we employed ANOVA (with the cancer type included in the model to account for its potential effect). We assessed the correlations of the mtDNA copy number with the patient’s age, overall survival and stage using Spearman’s rank correlation, cox model/log-rank test, and ANOVA, respectively. We log2-transformed the mtDNA copy number values when using ANOVA and t-test to conform to the normality assumption.

### Co-expression analysis

For each cancer type, we used the WGCNA package^33^ to build a weighted gene co-expression network that contains ~20,000 nodes (including both nuclear genes and mitochondrial genes). The key parameter, β, for a weighted network construction was optimized to maintain both the scale-free topology and sufficient node connectivity as recommended in the manual. In such a network, any two genes were connected and the edge weight was determined by the topology overlap measure provided in WGCNA. This measure considered not only the expression correlation between two partner genes but also how many ‘friends’ the two genes shared. The weights ranged from 0 to 1, which reflects the strength of the connection between the two genes. To identify mitochondria-related pathways, we performed GSEA^26^ on the basis of the full set of nuclear protein-coding genes, ranked on the basis of their weights of the edge connecting the mitochondrial genes, and detected significant pathways at FDR = 0.05. To construct the mitochondria-centric network, we focused on the top 500 neighboring genes that showed the strongest connections with the mitochondrial genes, with a minimum weight of 0.05. Among these neighboring genes, we detected the clinically actionable genes (defined as FDA-approved therapeutic targets and their relevant predictive markers^48^) in at least one of the cancer types we surveyed. We examined the correlations of mtDNA gene expression levels with mtDNA copy numbers using Spearman Rank correlations.

### TCMA data portal construction

We stored the pre-calculated mtDNA molecular data (including mtDNA mutation, nuclear transfer, copy number and expression) in a database of CouchDB. The Web interface was implemented by JavaScript; tables were visualized by DataTables; the co-expression network visualization was implemented by Cytoscape Web.

## Acknowledgements

We gratefully acknowledge contributions from the ICGC and especially the ICGC Pan-Cancer Analysis of Whole Genomes Group. This study was partially supported by the National Institutes of Health (U24CA143883, R01CA175486, U24CA209851 to H.L and the CCSG grant CA016672); a grant from the Cancer Prevention and Research Institute of Texas (RP140462 to H.L.); and the Lorraine Dell Program in Bioinformatics for Personalization of Cancer Medicine (to J.N.W.). This work was also supported by Institute for Information & communications Technology Promotion grant funded by the Korea government (MSIP) (No.B0101-15-0104, The Development of Supercomputing System for the Genome Analysis) and by a grant of Korea Health Technology R&D Project through the Korea Health Industry Development Institute (KHIDI), funded by the Ministry of Health & Welfare, Republic of Korea (grant no: HI14C0072 to K.P. and HI16C2387 toY.S.J.). We also thank ETRI (Electronics and Telecommunications Research Institute, Korea) for the commitment to the ICGC-PCAWG projects, the MD Anderson high-performance computing core facility for computing, and LeeAnn Chastain for editorial assistance.

